# Whole-bulb LSFEM enables automated reconstruction and quantitative spatial mapping of thousands of olfactory glomeruli

**DOI:** 10.64898/2026.11.16.766961

**Authors:** Juan Edo Rodriguez-Gatica, Joyce Imaculate Jayakumar, Sharon Innocent Gnanasekar, Ali Mohammadi, Dominique Roos, Tobias Ackels, Markus Rothermel, Martin K. Schwarz

**Affiliations:** Institute for Experimental Epileptology and Cognition Research, University of Bonn Medical Faculty, Venusberg-Campus1, 53127 Bonn; Institute of Physiology, RG Neurophysiology and Optogenetics, Medical Faculty, Otto-von-Guericke-University, 39120 Magdeburg

**Keywords:** olfactory bulb, glomerulus, light-sheet microscopy, expansion microscopy, SegFormer, semantic segmentation, three-dimensional connected components analysis

## Abstract

The glomerular layer of the olfactory bulb (OB) transforms chemical information into the first spatially organized neural representation of odor identity. Yet its three-dimensional architecture remains difficult to quantify at whole-bulb scale. Here, we establish an end-to-end framework integrating light-sheet fluorescence expansion microscopy (LSFEM) with automated reconstruction of thousands of individual glomeruli across intact mouse OBs. LSFEM preserves whole-bulb architecture and supports genetically encoded fluorescence and immunolabelling, providing continuous multichannel input for automated analysis. SegFormer-based segmentation of 2D optical sections was followed by morphology-guided separation and three-dimensional object reconstruction. Across 12 intact bulbs, the workflow yielded a median of 2,677 glomeruli per bulb, approaching the 2,851 reported by complete three-dimensional reconstruction. Object-level maps further revealed that glomeruli spanning a broad range of volumes were intermingled throughout the bulb without evident size-defined anatomical domains. Because each reconstructed glomerulus retains a three-dimensional centroid coordinate, the resulting object map can additionally be used as a specimen-specific spatial reference framework, allowing distances and distributions to be expressed relative to defined glomerular or anatomical positions within the intact OB. Together, whole-bulb LSFEM and automated reconstruction provide a scalable framework for quantitative three-dimensional mapping of glomerular number, morphology, and spatial organization, and the relative positioning of additional anatomical signals within the glomerular architecture.

## Introduction

The mammalian olfactory bulb (OB) performs an early transformation of sensory information, converting a high-dimensional chemical input space into structured neural representations that guide behavior. Olfactory sensory neurons expressing the same odorant receptor converge onto a small number of glomeruli (Mori and Sakano, 2011), organizing peripheral input into receptor-defined channels relayed to mitral and tufted cells (e.g. Nagayama et al., 2014). Glomeruli therefore provide the anatomical basis for the combinatorial coding of odor identity and define the spatial units within which olfactory processing is organized (Buck and Axel, 1991; Mombaerts et al., 1996; Egger and Sachse, 2026; Sireci and Rothermel, 2026). Their arrangement is functionally important because glomeruli form discrete sensory channels whose activity is shaped by respiration as well as by local and interglomerular circuit interactions, contributing to the transformation and integration of odor information across channels (e.g. Wachowiak, 2011; Mori and Sakano, 2021; Ackels et al., 2020; Tootoonian et al., 2026; Strauch et al., 2026).

Despite their central role, the total number and three-dimensional morphology of glomeruli have been difficult to determine. Classical estimates based on sparse histological sections ranged from approximately 1,800 to 3,600 glomeruli per mouse OB and depended heavily on assumptions about object diameter and shape (Royet et al., 1988; Richard et al., 2010; Weng et al., 2025). Complete serial two-photon tomography and manual three-dimensional segmentation recently yielded an empirical median of 2,851 glomeruli in adult C57BL/6J bulbs (Weng et al., 2025). That study further demonstrated that glomeruli are not spherical but rather rounded, elongated, and irregular, described as tuberiform, with a median effective diameter of 77.5 µm and a median sphericity of 0.56 in adult animals (Weng et al., 2025). These observations highlight two related analytical challenges: individual glomeruli cannot be reliably represented by a single two-dimensional diameter, and densely apposed, non-spherical objects are difficult to separate automatically.

Volumetric fluorescence imaging provides a route to resolve these limitations, but it shifts the bottleneck from image acquisition to image analysis. Light-sheet fluorescence microscopy rapidly captures large transparent samples with limited out-of-focus excitation, while expansion microscopy physically enlarges tissue to improve effective resolution (Bürgers et al., 2019; Chen et al., 2015; Düring et al., 2019; Mascheroni et al., 2020; Pesce et al., 2024; Melhem et al., 2026). By combining these approaches, light-sheet fluorescence expansion microscopy (LSFEM) enables high-resolution imaging across large neural volumes and is particularly well suited to the continuous architecture of the glomerular layer. Importantly, LSFEM maintains the sample’s continuous three-dimensional organization, enabling imaging of the intact OB without physical sectioning. Using a controlled two-fold expansion (Rodriguez-Gatica et al., 2022), the complete OB can be visualized while retaining information on its cellular and structural organization. The approach is compatible with nuclear staining, preservation of genetically encoded fluorescent proteins, and immunolabeling with conventional antibodies, providing a basis for relating glomerular organization to other cellular and anatomical features throughout the intact OB (Figure S1) (Melhem et al., 2026; Rodriguez-Gatica et al., 2022).

A single OB volume, however, contains hundreds of high-resolution sections and thousands of glomeruli to be identified and quantified. Manual delineation is therefore laborious. Complete manual segmentation of an entire bulb required an average of approximately 90 hours per bulb (Weng et al., 2025). Automated approaches have been developed for smaller or more isolated chemosensory structures within the accessory OB (Bahreini Jangjoo et al., 2021), but robust bulb-wide analysis remains challenging.

The rich three-dimensional information provided by LSFEM therefore requires an analysis strategy that can efficiently extract glomerular architecture across whole-bulb datasets. Deep learning offers an opportunity to replace repeated manual delineation with reproducible image segmentation. Convolutional architectures such as U-Net are widely used in biomedical imaging (Ronneberger et al., 2015), whereas hierarchical transformers can integrate local image features with long-range spatial context. SegFormer couples a hierarchical multiscale transformer encoder with a lightweight multilayer perceptron decoder, enabling dense semantic segmentation without relying on explicit positional encodings (Xie et al., 2021). In OB images, however, semantic foreground prediction is not sufficient for object-level analysis because adjacent glomeruli are often merged into contiguous regions. A complete computational workflow therefore requires three sequential components: learned foreground segmentation, morphology-based instance separation of clustered glomeruli, and three-dimensional association of corresponding glomerular candidates across consecutive optical sections.

Here, we establish an end-to-end framework that converts intact whole-bulb LSFEM datasets into quantitative three-dimensional maps of thousands of individual glomeruli. We manually annotated optical LSFEM sections to train a SegFormer model, then automatically separated predicted glomerular foreground into individual candidates and linked these candidates across the z-axis to reconstruct three-dimensional objects. Applied to 15 OB volumes, the workflow enabled bulb-wide reconstruction and quantification of individual glomeruli. Across the 12 intact datasets, automated analysis yielded a median of approximately 2,677 glomeruli per bulb, closely approaching the median of 2,851 obtained by recent complete manual three-dimensional reconstruction (Weng et al., 2025). Beyond counting, the resulting object-level maps retained information about glomerular morphology and spatial position, revealing that glomeruli spanning a broad range of volumes were intermingled throughout the bulb rather than segregated into obvious size-defined domains. Collectively, our results establish a scalable route from intact whole bulb imaging to the automated reconstruction and quantitative analysis of thousands of individual glomeruli within their native three-dimensional organization.

## Materials and Methods

### Biological metadata

OB tissue was obtained from C57BL/6J mice at postnatal ages P7–P212. Datasets included only male mice. The dataset therefore comprised both juvenile (P7 and P14) and adult (P210–P212) animals. Strain, age, and sex were recorded for each dataset and are reported in the accompanying dataset metadata table. Red fluorescent cells were imaged from Tbet-Cre:Ai14 mice, generated by crossing the Tbet-Cre driver line (Haddad et al., 2013; JAX stock #024507) with the Ai14 tdTomato reporter line (Madisen et al., 2010; JAX stock #007914). Cre-dependent tdTomato expression labelled mitral and tufted cell bodies and dendritic processes within the OB.

| Dataset No. | Age | Sex | Line/Strain |
|---|---|---|---|
| 1 | P210 | ♂ | 1 - C57BL/6J |
| 2 | P210 | ♂ | 1 - C57BL/6J |
| 3 | P211 | ♂ | 1 - C57BL/6J |
| 4 | P211 | ♂ | 1 - C57BL/6J |
| 5 | P212 | ♂ | 1 - C57BL/6J |
| 6 | P211 | ♂ | 1 - C57BL/6J |
| 7 | P210 | ♂ | 1 - C57BL/6J |
| 8 | P212 | ♂ | 1 - C57BL/6J |
| 9 | P7 | ♂ | Thy1-GFP-M |
| 10 | P7 | ♂ | Thy1-GFP-M |
| 11 | P14 | ♂ | Thy1-GFP-M |
| 12 | P14 | ♂ | Thy1-GFP-M |
| 13 | P210 | ♂ | Tbet-Cre:Ai14 tdTomato |
| 14 | P7 | ♂ | Thy1-GFP-M |
| 15 | P7 | ♂ | Thy1-GFP-M |

### Sample preparation and LSFEM microscopy

#### Sample preparation

Whole mouse olfactory bulbs (OBs) were processed using a light-sheet fluorescence expansion microscopy (LSFEM) protocol adapted from Rodriguez-Gatica et al. (2022). Briefly, whole OBs were incubated with 2 mM methacrylic acid N-hydroxysuccinimide ester (MA-NHS) for 48 h at room temperature (RT) with gentle agitation. Samples were subsequently washed three times in PBS for 45 min each and incubated for 48 h at 4°C with agitation in monomer solution containing 8.6% sodium acrylate, 2.5% acrylamide, 0.15% N,N′-methylenebisacrylamide, and 11.7% NaCl in 1× PBS.

For polymerization, fresh monomer solution was supplemented with 0.01% 4-hydroxy-TEMPO, 0.2% TEMED, and 0.2% ammonium persulfate. Samples were initially maintained on ice to prevent premature polymerization, incubated in gelling solution for 5 min at 4°C with agitation, transferred to a gelling chamber, and polymerized overnight at 4°C, followed by an additional 2 h incubation at 37°C. The resulting gels were digested at 37°C in digestion buffer containing 50 mM Tris, 1 mM EDTA, 0.5% Triton X-100, 0.8 M guanidine HCl, and 16 U/mL proteinase K (pH 8.0). Digestion buffer was replaced every 24–48 h as needed, and complete digestion of whole OBs typically required about 3 days. Following digestion, samples were washed three times with PBS and stored in PBS. Under these conditions, samples showed an approximately two-fold linear expansion relative to their original dimensions.

#### Immunolabeling

Following digestion, preserved tdTomato fluorescence was amplified by immunolabeling. Samples were blocked overnight at RT with agitation in blocking buffer containing 5% normal donkey serum, 0.1% Triton X-100, and 0.02% sodium azide in 1× PBS. Samples were then incubated for 48-72 h at 4°C with goat anti-tdTomato primary antibody (1:500; Sicgen, AB8181) diluted in blocking buffer. After three 40-min washes in blocking buffer at RT, samples were incubated for 48-72 h at 4°C with donkey anti-goat Alexa Fluor 555 secondary antibody (1:400; Invitrogen, A21432). Nuclei were counterstained with Hoechst 33342 (Invitrogen, H3570; 2.5 µg/mL).

#### Light-sheet fluorescence microscopy

Expanded whole OBs were imaged using an UltraMicroscope Blaze light-sheet fluorescence microscope (LaVision BioTec/Miltenyi Biotec). Digested samples were immobilized on a coverslip using poly-L-lysine, mounted in a custom sample holder, and immersed in imaging buffer consisting of 1× PBS supplemented with 0.02% sodium azide.

For whole-bulb mesoscopic imaging, fluorescence excitation was provided by fiber-coupled 405-, 488-, 561-, and 638-nm laser lines using dual-sided illumination. The light-sheet beam waist was adjusted to a 1/e² diameter of approximately 9 µm. Fluorescence was detected using a MI PLAN 4×/0.35 NA objective (15-mm working distance; LaVision BioTec/Miltenyi Biotec) with a water-dipping cap and an sCMOS camera (2048 × 2048 pixels; 6.5-µm pixel size) operated in rolling-shutter mode. This configuration provided an effective field of view of approximately 3.33 × 3.33 mm². Where necessary, multiple image volumes were acquired to encompass the complete expanded OB.

#### LSFEM data processing

Raw image data acquired with the UltraMicroscope Blaze were first converted to Imaris format (.ims) using Imaris Converter. Multiple three-dimensional image volumes were subsequently stitched to reconstruct the complete OB using Imaris Stitcher (Imaris, version 10.0.1; Bitplane Inc., Zurich, Switzerland). Three-dimensional visualization of the reconstructed datasets was performed using the Surpass view in Imaris.

For subsequent image analysis, the stitched “.ims” datasets were opened in Fiji/ImageJ using the Bio-Formats Importer plugin (Schindelin et al., 2012). Individual fluorescence channels were separated, and the Hoechst 33342 nuclear-staining channel was exported as a TIFF image stack. These TIFF image stacks were subsequently used as input for the image preprocessing and SegFormer-based glomerular segmentation pipeline described below.

Image processing, reconstruction, and analysis were performed using two dedicated high-performance workstations. Workstation 1 was equipped with an Intel® Xeon® Gold 6128 CPU at 3.40 GHz (6 cores/12 logical processors), 128 GB DDR4 RAM, and an NVIDIA GeForce GTX 1080 Ti GPU, and operated under Windows 10. Workstation 2 was equipped with two Intel® Xeon® Platinum 8160 CPUs at 2.10 GHz (48 cores/96 logical processors in total), 512 GB DDR4 RAM, and an NVIDIA Quadro P5000 GPU, operated under Windows 11 Pro.

#### Image data and study design

LSFEM image stacks of expanded mouse OBs generated as described above were used for development and evaluation of the automated glomerular segmentation and three-dimensional quantification pipeline. The source datasets comprised 16-bit TIFF images with dimensions up to 6118 × 3806 pixels, typically 300–750 sections per bulb, and a z-step of 10 µm. The imaged volume per dataset was approximately 20–30 mm³. Fifteen bulb volumes were processed for final three-dimensional counting. The model-development workflow incorporated six independent OB datasets. Across these datasets, a total of 290 optical sections were manually annotated, of which 207 were used for model training and 83 for validation.

#### Ground-truth annotation and data split

Glomerular regions were manually delineated using the MATLAB Image Labeler toolbox. Annotations were exported as binary masks with foreground glomeruli assigned a value of 1 and background assigned a value of 0. Two trained annotators generated manual annotations independently. A subset of the annotations was repeated independently by the second annotator to assess consistency, and additional sections were subsequently annotated to expand the dataset. The resulting masks were reviewed for consistency in the delineation of glomerular regions. In total, 290 manually annotated optical sections from six independent OB datasets were included for model development and evaluation. For each dataset, approximately 70–80% of the annotated sections were assigned to the training set and the remaining 20–30% to the validation set, resulting in 207 training images and 83 validation images overall. Across the 80 training sections for which glomerular profile counts were quantified, the mean number of manually delineated glomerular profiles was 76 ± 60 per section. Across all 207 training sections, glomerular foreground represented 1.2 ± 0.9% of pixels per section (mean ± SD), highlighting the pronounced class imbalance between glomerular foreground and background. Individual optical sections were assigned exclusively to either the training or validation subset and were never used in both. Because the split was performed at the section level within each dataset, however, different sections originating from the same OB could occur in both subsets.

#### Image preprocessing

Raw images were converted to 8-bit grayscale TIFF files for model training and inference. The images were then rescaled to 1/3 of their original size while maintaining the aspect ratio. Brightness and contrast were adjusted uniformly across each stack using Fiji/ImageJ, corresponding to a linear intensity rescaling. Images were sharpened using a Laplacian edge-enhancement operation. Finally, an iterative histogram-based thresholding method was applied across the stack to convert grayscale images to binary images and set uniform intensity. The same preprocessing settings were applied throughout the stack to reduce variation.

#### SegFormer training and inference

A transformer-based semantic segmentation approach was developed for automated detection of glomeruli in LSFEM images of the OB. The segmentation framework was adapted from a publicly available SegFormer-based medical image segmentation workflow demonstrated on the UW-Madison GI Tract Image Segmentation dataset and implemented using the Hugging Face Transformers and PyTorch frameworks. SegFormer combines a hierarchical transformer encoder with a lightweight all-MLP decoder, eliminating the need for complex positional encodings while efficiently capturing local and global multi-scale features. This design delivers superior segmentation performance and robustness compared to traditional CNN architectures while maintaining a significantly more efficient and computationally lightweight footprint. The original workflow was modified for OB glomerular segmentation by replacing the dataset-specific preprocessing pipeline, annotations, and segmentation classes with glomeruli-specific image– mask pairs generated from LSFEM datasets.

The segmentation model used the SegFormer-B0 architecture. The network was initialized using publicly available NVIDIA SegFormer-B0 weights pretrained on the ADE20K semantic segmentation dataset and further fine-tuned on manually annotated LSFEM images. The final segmentation head was modified to perform binary semantic segmentation, assigning each pixel to either background or glomerular foreground.

The dataset was divided into training and validation subsets, and the model was optimized using a Dice-based loss function to improve segmentation accuracy for small, irregularly shaped glomerular structures. Model parameters were updated using the Adam optimizer with a learning rate of for 40 epochs with a batch size of 2 at a learning rate of 1×10^−4^.

Following training, the optimized model was applied to previously unseen LSFEM images to generate pixel-level glomerular-foreground probability maps. No learning-rate scheduler, early stopping, or data augmentation was used in the final configuration.

#### Separation of merged glomeruli

SegFormer produced foreground masks in which adjacent glomeruli could remain connected. To obtain individual glomerular candidates from connected foreground regions, masks were smoothed with a Gaussian filter (σ = 0.5), binarized, and cleared of components smaller than 5 pixels. Within each connected foreground region, an iterative H-maxima transform was used to identify stable internal peaks. The height parameter began at 0.01 and increased in increments of 0.005 until the number of maxima remained stable for five iterations. For each retained maximum, an external Euclidean distance transform was calculated and normalized by the mean marker height. The minimum of these weighted transforms was Gaussian-smoothed and used as the topographic surface for marker-controlled watershed segmentation, following the general strategy of Portuondo-Mallet and colleagues (Portuondo-Mallet et al., 2022). Watershed labels were converted back to binary masks for subsequent volume assembly.

#### Three-dimensional reconstruction and quantification

Three-dimensional connected pixels were identified with scikit-image’s three-dimensional connected component analysis tool. Using connectivity = 2 in scikit-image corresponds to 18-connectivity in three dimensions, such that voxels sharing a face or edge are connected while corner-only contacts remain separate. This intermediate connectivity was selected to reduce fragmentation of obliquely continuous structures without merging objects through isolated diagonal contacts. Each three-dimensional connected component was assigned a unique identifier and treated as a reconstructed glomerular object for subsequent quantification. Region properties, including centroid coordinates and voxel count, were exported as comma-separated values. Volumes were visualized by direct volume rendering using the napari framework.

#### Performance evaluation

Pixel-level predictions were compared with manual masks on the held-out annotated subset. Reported measures included accuracy, specificity, precision, recall, Dice/F1 score, Matthews correlation coefficient (MCC), and a dataset-level receiver-operating-characteristic area under the curve (ROC AUC). Metrics were calculated independently for each validation section and are reported as mean ± SD across sections. The behavior of the three-dimensional linking procedure was additionally evaluated under idealized axial continuity using a synthetic repeated-slice control (Supplementary Figure S2).

#### External anatomical benchmark

Automated counts were compared descriptively with the empirical three-dimensional census of Weng and colleagues, who manually segmented 33,137 VGLUT2-positive glomeruli in 12 mouse OBs and reported a median of 2,851 glomeruli per adult PD56 bulb (Weng et al., 2025). This comparison was used solely as an anatomical scale reference rather than as a formal accuracy or biological comparison because the datasets differed in imaging modality, tissue preparation, labeling, spatial sampling, animal age, and strain. In particular, the present dataset included both juvenile (P7 and P14) and adult (P210) animals, whereas the Weng et al. dataset comprised P56 adult mice.

## Results

### LSFEM preserves whole-bulb three-dimensional architecture and molecular information

We first established that LSFEM enables continuous visualization of cellular and anatomical organization throughout the intact mouse OB while preserving its three-dimensional architecture without physical sectioning. Following two-fold expansion, whole OB samples retained their overall morphology and could be imaged throughout their complete volume by light-sheet fluorescence microscopy (Figure 1A, B). Nuclear staining with Hoechst 33342 revealed the cellular organization across the bulb and delineated glomeruli as cell-poor regions, whereas preserved tdTomato fluorescence visualized the dendritic arborization of genetically labeled mitral/tufted cells. At higher magnification, tdTomato-positive processes could be seen extending into these Hoechst-defined cell-poor regions, providing complementary anatomical information supporting their identification as individual glomeruli (Figure 1B–C; Figure S1A–C) (Haddad et al., 2013). The combination of these complementary signals therefore allowed glomerular structures to be identified within their preserved three-dimensional anatomical context. Orthogonal views further demonstrated that cellular and glomerular organization could be examined across multiple spatial planes without disrupting their relative three-dimensional organization (Figure 1C).

**Figure 1.**
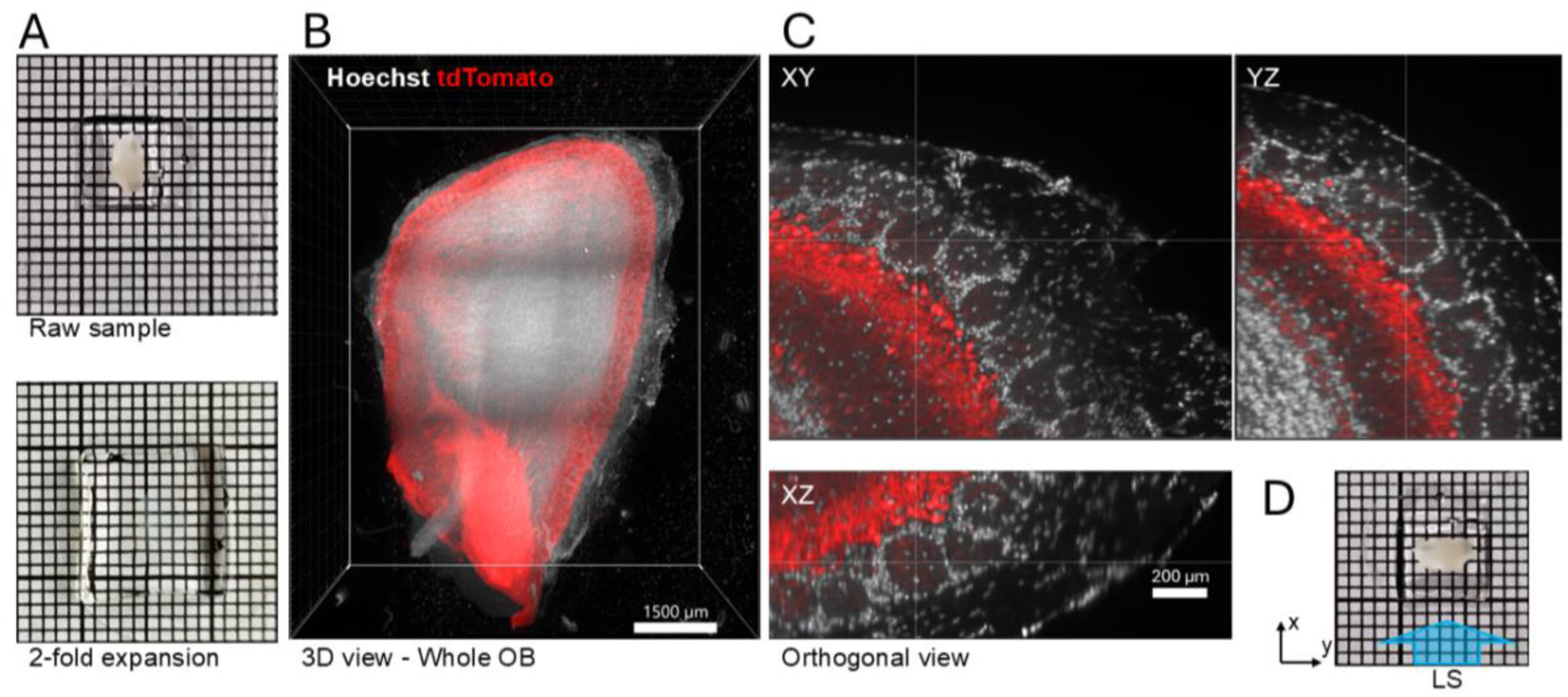
LSFEM enables three-dimensional visualization of the intact expanded mouse OB. (A) Representative OB before and after approximately 2-fold expansion (left). (B) Three-dimensional rendering of the complete expanded OB acquired by light-sheet fluorescence microscopy, showing Hoechst nuclear staining (gray) and preserved tdTomato fluorescence in mitral/tufted cells (red). (C) Orthogonal views of the marked region of interest in B, illustrate the distribution of nuclear and tdTomato signals within the intact three-dimensional volume (right). (D) Orientation of the expanded sample relative to the light-sheet (LS) illumination during imaging.

Importantly, the expanded whole-bulb preparations also remained compatible with conventional immunolabeling. Neurofilament-H staining produced robust structural labeling throughout the expanded tissue and could be visualized together with Hoechst and preserved tdTomato fluorescence within the same anatomical region (Figure S1D), demonstrating that additional molecular and cellular information can be incorporated into the same intact three-dimensional preparation. Thus, LSFEM is not restricted to visualizing glomeruli alone but provides a multichannel whole-organ imaging framework in which glomerular architecture can be reconstructed together with genetically encoded and immunolabeled cellular features. This combination of intact whole-bulb reconstruction with genetically encoded fluorescence and post-expansion immunolabeling provides a direct route toward quantitative three-dimensional mapping of glomeruli in relation to defined neuronal populations and anatomical structures across the complete OB.

### Manual annotation generated ground-truth masks of the glomerular layer

We next asked whether the LSFEM volumes contained a sufficiently consistent anatomical signal for automated segmentation. In individual optical sections of the Hoechst channel, glomeruli could be reliably identified along the peripheral glomerular layer based on their characteristic organization and contrast relative to the surrounding tissue (Figure 2A). We manually delineated individual glomeruli to generate ground-truth segmentation masks, assigning each annotated glomerular region to the foreground (Figure 2B). Overlaying the manually generated annotations onto the corresponding LSFEM images confirmed their spatial correspondence with the glomerular structures identified in the Hoechst channel (Figure 2C). In total, we generated 290 annotated slices from six datasets for model development. Across the annotated sections, the mean number of manually delineated glomerular profiles was 76 ± 60 per section (mean ± SD). These manually annotated images provided the reference masks used to train and evaluate the SegFormer-based segmentation model.

**Figure 2.**
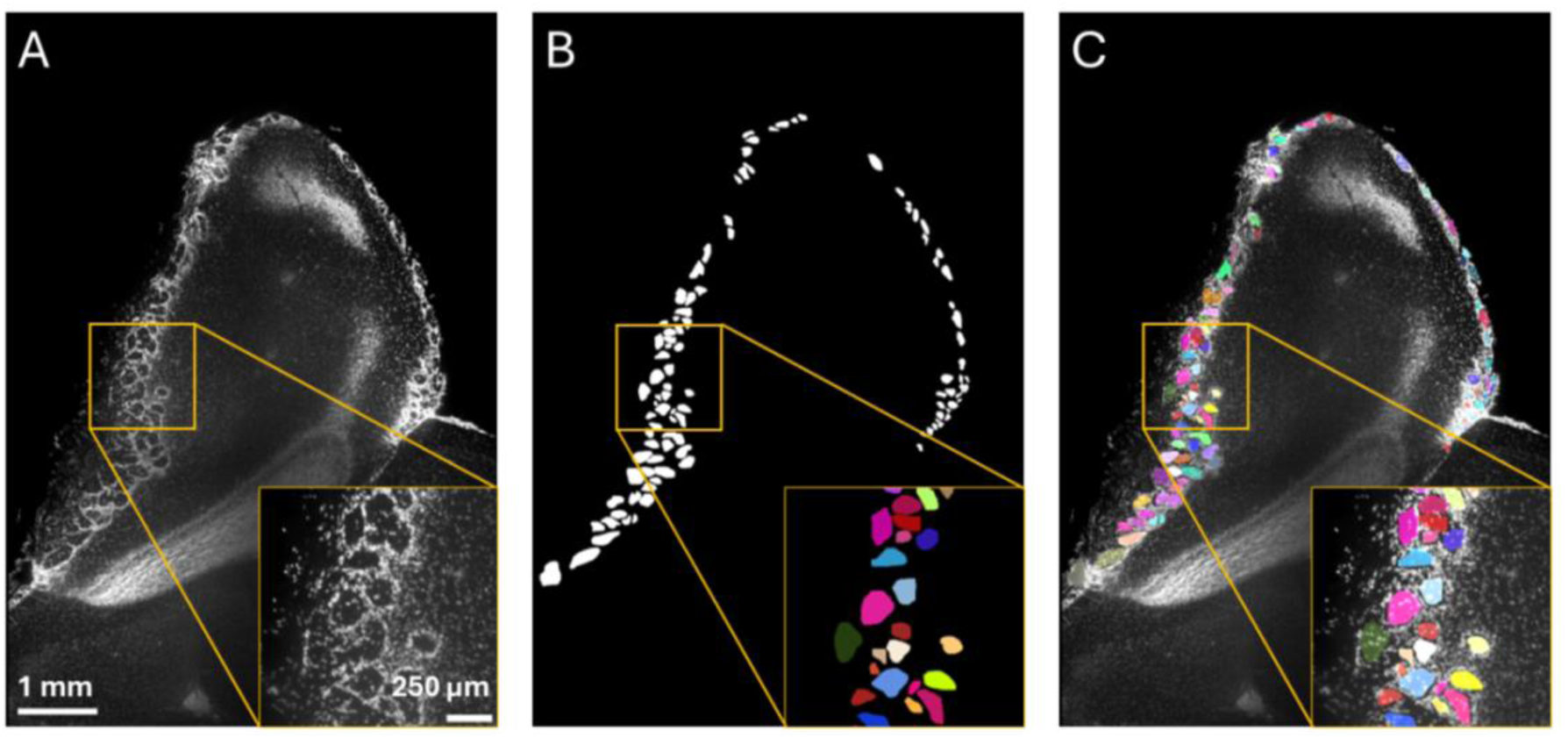
Manual annotation of OB glomeruli in optical cross sections of LSFEM-generated images. (A) Representative optical section of an expanded mouse OB imaged by LSFEM using Hoechst nuclear staining. The boxed region indicates a representative region of interest (ROI) within the glomerular layer, shown at higher magnification in the inset. (B) Corresponding manually generated binary ground-truth mask of the glomerular layer. Individual glomeruli in the enlarged ROI are displayed using arbitrary colors for visualization purposes only; binary masks were used for automated segmentation. (C) Overlay of the manual annotations onto the corresponding LSFEM image, demonstrating the spatial correspondence between the annotated regions and glomerular structures.

### SegFormer consistently identified the glomerular layer across different optical sections

To automate glomerular layer identification, we used SegFormer, a transformer-based semantic segmentation architecture that combines hierarchical multiscale feature extraction with contextual information and a lightweight multilayer perceptron (MLP) decoder to assign individual pixels to predefined classes (Xie et al., 2021) (Figure 3).

**Figure 3.**
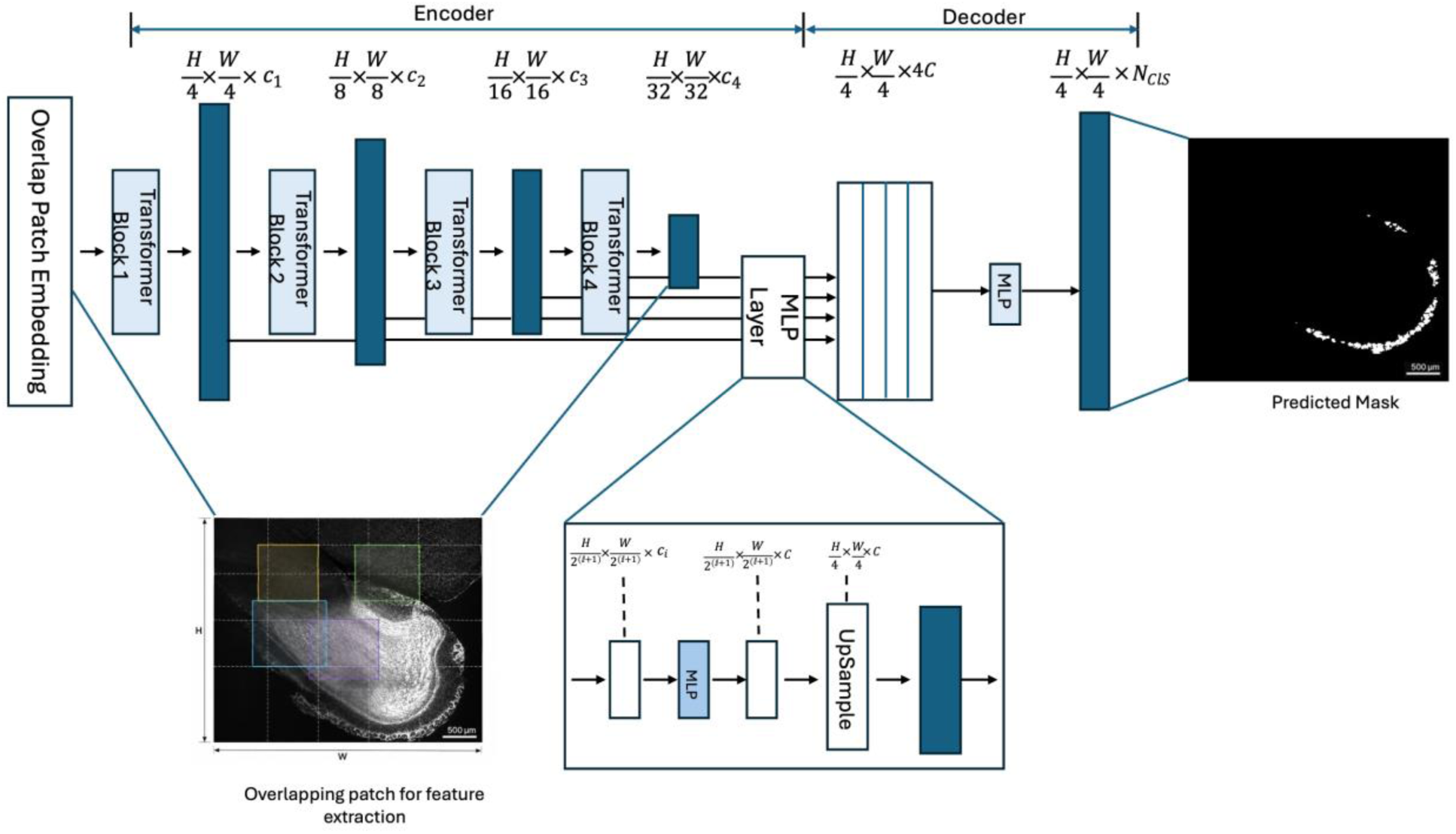
Architecture of the SegFormer-based segmentation network tailored for processing LSFEM image sections of the OB. The framework comprises a hierarchical transformer encoder with overlapping patch embeddings and four progressive downsampling blocks producing multi-scale feature maps with channel dimensions *c*_1_ through *c*_4_, paired with a lightweight MLP decoder that fuses these multi-level representations to generate the final glomerular foreground mask *Ncls*.

In our implementation, we trained SegFormer on preprocessed Hoechst optical sections and corresponding manual annotations, which generated binary glomerular-foreground masks representing the spatial distribution of the glomerular layer. Importantly, at this stage the network performs semantic segmentation and therefore identifies glomerular foreground rather than assigning identities to individual glomeruli. When applied to previously unseen images, the model generated predicted masks that followed the peripheral distribution of glomeruli across the OB (Figure 4). This spatial correspondence was maintained across different positions within the same OB volume, as illustrated by slices 580 and 627 of a 759-slice z-stack, despite changes in tissue size, shape, and appearance across the stack (Figure 4A–C). Overlaying the predicted masks onto the corresponding Hoechst images confirmed that the segmented regions were accurately localized to the glomerular layer (Figure 4C). Although the model successfully localized the glomerular layer, semantic segmentation did not yet accurately resolve individual glomeruli. We observed that, in densely packed regions, adjacent glomeruli could be represented as a single connected foreground region. Thus, while SegFormer provided a robust representation of the glomerular layer’s location and continuity, an additional object-separation step was required before individual glomeruli could be identified and subsequently quantified in three dimensions.

**Figure 4.**
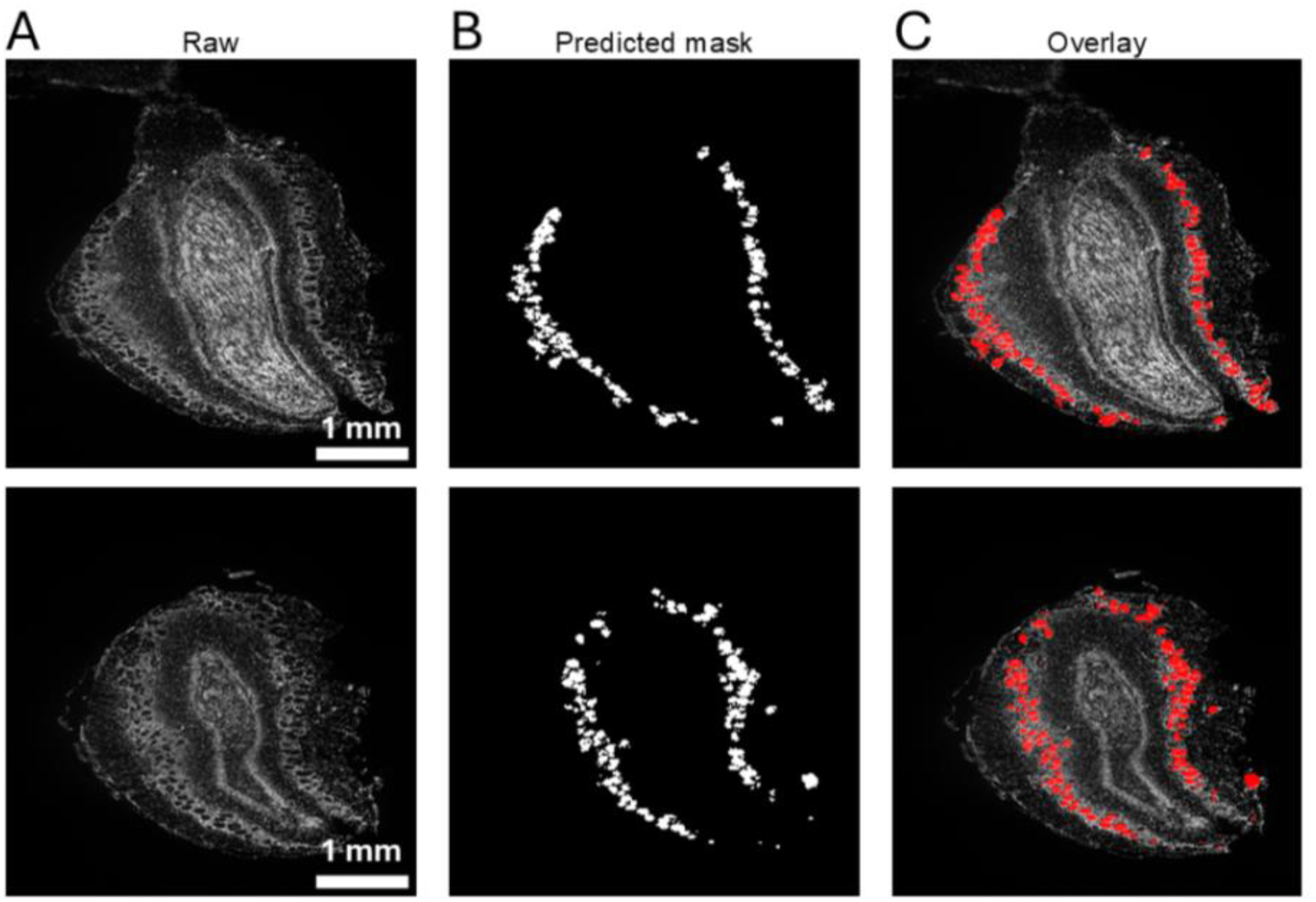
SegFormer identification of the glomerular layer across different optical sections. (A) Representative LSFEM images corresponding to slices 580 (top) and 627 (bottom) of a 759-slice z-stack acquired from the same expanded mouse OB, visualized using the Hoechst channel. (B) Corresponding binary masks predicted by the fine-tuned SegFormer model. (C) Overlay of the predicted masks (red) onto the corresponding LSFEM images, demonstrating that the predicted regions predominantly follow the peripheral glomerular layer at different positions within the z-stack.

### Morphological post-processing separated merged predictions into individual glomerular candidates

To enable accurate object-level analysis, we applied a morphology-based post-processing procedure to separate the merged glomerular predictions. Stable internal markers were identified within connected foreground regions using iterative H-maxima detection. These markers, together with a weighted Euclidean distance transform, were subsequently used to guide watershed segmentation (Portuondo-Mallet et al., 2022) and partition connected regions into individual glomerular objects (for a detailed description, please see Materials and Methods). As illustrated in Figure 5, foreground regions represented as connected structures in the SegFormer prediction could then be resolved into spatially distinct objects (Figure 5A, B lower panels). The resulting separated objects were assigned individual labels, displayed using arbitrary colors for visualization (Figure 5B), providing the object-level representation required for subsequent three-dimensional reconstruction and quantification.

**Figure 5.**
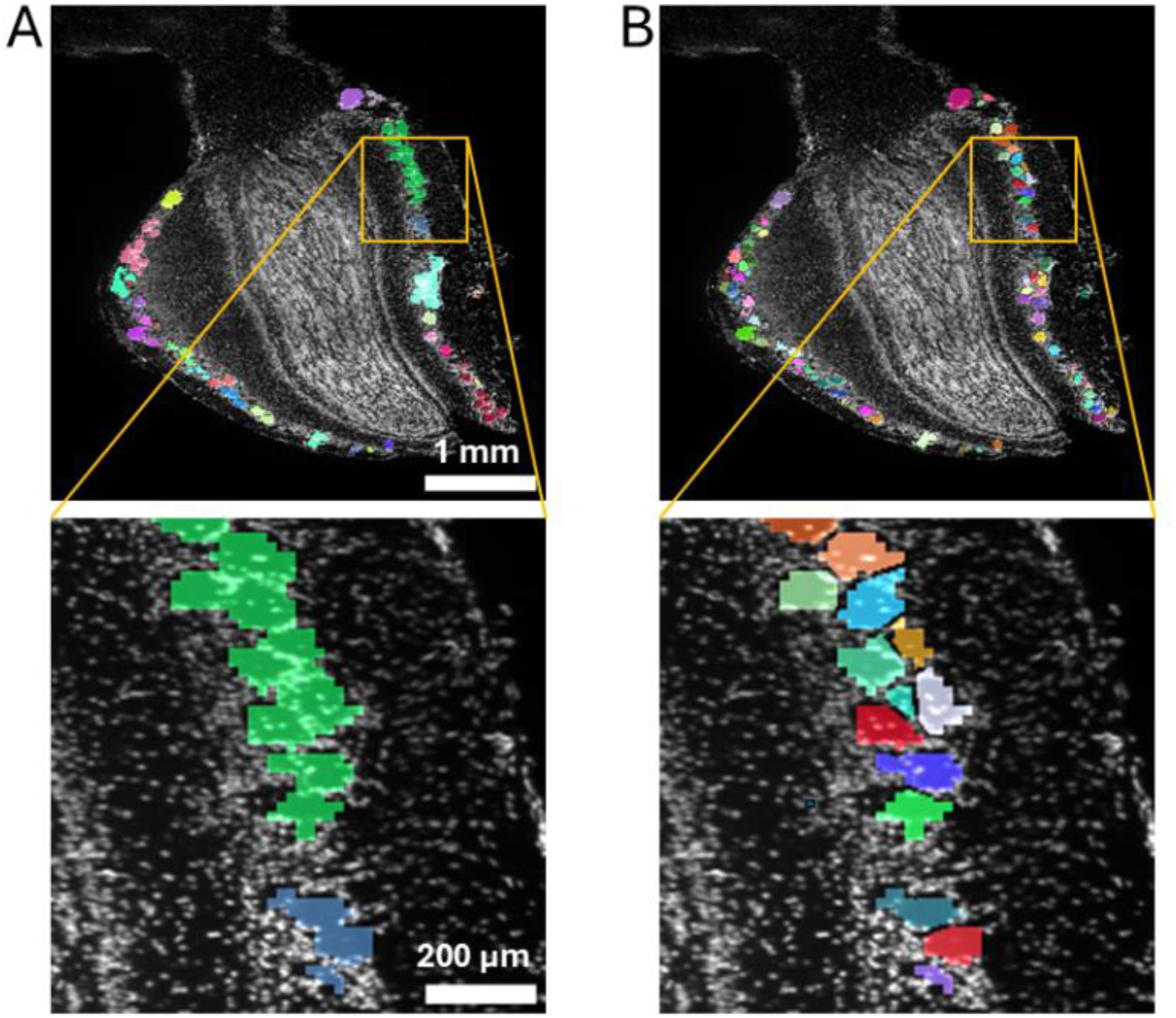
Morphological post-processing separates connected SegFormer predictions into individual glomeruli. (A) SegFormer prediction overlaid on the corresponding Hoechst LSFEM image. Predicted glomerular regions (connected regions) in the SegFormer prediction are displayed using arbitrary colors for visualization only; the SegFormer output itself was a binary semantic mask. The boxed region and enlarged view illustrate an area where multiple closely apposed glomeruli remain connected within the predicted mask. (B) Corresponding result following morphology-based post-processing and watershed separation. Individual separated glomerular candidates are displayed using arbitrary colors. The enlarged region illustrates the separation of connected predictions into discrete glomerular candidates suitable for subsequent three-dimensional reconstruction and analysis.

### The combined workflow retained foreground information despite strong class imbalance

We next quantified how reliably the SegFormer predictions reproduced the manual foreground annotations. On independent test slices, pixel-wise accuracy reached 98.71% and specificity 99.38% (Table 1; Figure 5A), indicating that the workflow reliably excluded the extensive non-glomerular background. However, because background pixels strongly outnumbered glomerular pixels, these values alone provide limited information about foreground segmentation. Foreground precision was 74.18%, indicating that approximately three-quarters of pixels classified as glomerular corresponded to the manual annotations, while a recall of 70.46% showed that the workflow recovered approximately seven out of ten manually annotated glomerular pixels. The resulting Dice/F1 score of 72.65% reflects the balance between these two measures. Overall classification performance was further supported by a Matthews Correlation Coefficient (MCC) of 0.895 and a dataset-level ROC AUC of 0.849. The slightly higher precision than recall is consistent with a conservative segmentation strategy, in which glomerular regions were reliably identified while some foreground signal, particularly at low-contrast boundaries or between densely apposed structures, remained unassigned. Together, these results demonstrate robust recovery of glomerular foreground despite the pronounced pixel-class imbalance.

**Table 1.** Pixel-level performance reported for the held-out annotated slices.

| **Metric** | **Reported value** |
|---|---|
| Accuracy | 98.71% |
| Specificity | 99.38% |
| Precision | 74.18% |
| Recall | 70.46% |
| Dice/F1 | 72.65% |
| MCC | 0.895 |
| ROC AUC | 0.84884 |

### Manual three-dimensional segmentation validated automated glomerular reconstruction

As a complementary technical control, we first confirmed that repeated representations of the same glomerular profiles across consecutive optical sections were linked into continuous three-dimensional objects rather than counted independently (Supplementary Figure S2). To validate three-dimensional reconstruction under biologically realistic conditions, we compared the automated workflow with manual segmentation of glomeruli within a selected three-dimensional region of interest from an LSFEM OB dataset (Figure 6A). Individual glomeruli were manually identified and followed across consecutive optical sections to generate a reference three-dimensional reconstruction (Figure 6B). The same region was independently processed using the automated SegFormer-based segmentation, morphological separation, and three-dimensional linking workflow (Figure 6C). The automated reconstruction closely reproduced the overall number and spatial organization of manually identified glomerular objects. Manual reconstruction identified 28 individual glomeruli, whereas the automated workflow detected 27 objects, corresponding to a difference of one object. Thus, under biologically realistic imaging conditions, the automated pipeline closely approximated manual three-dimensional reconstruction.

**Figure 6.**
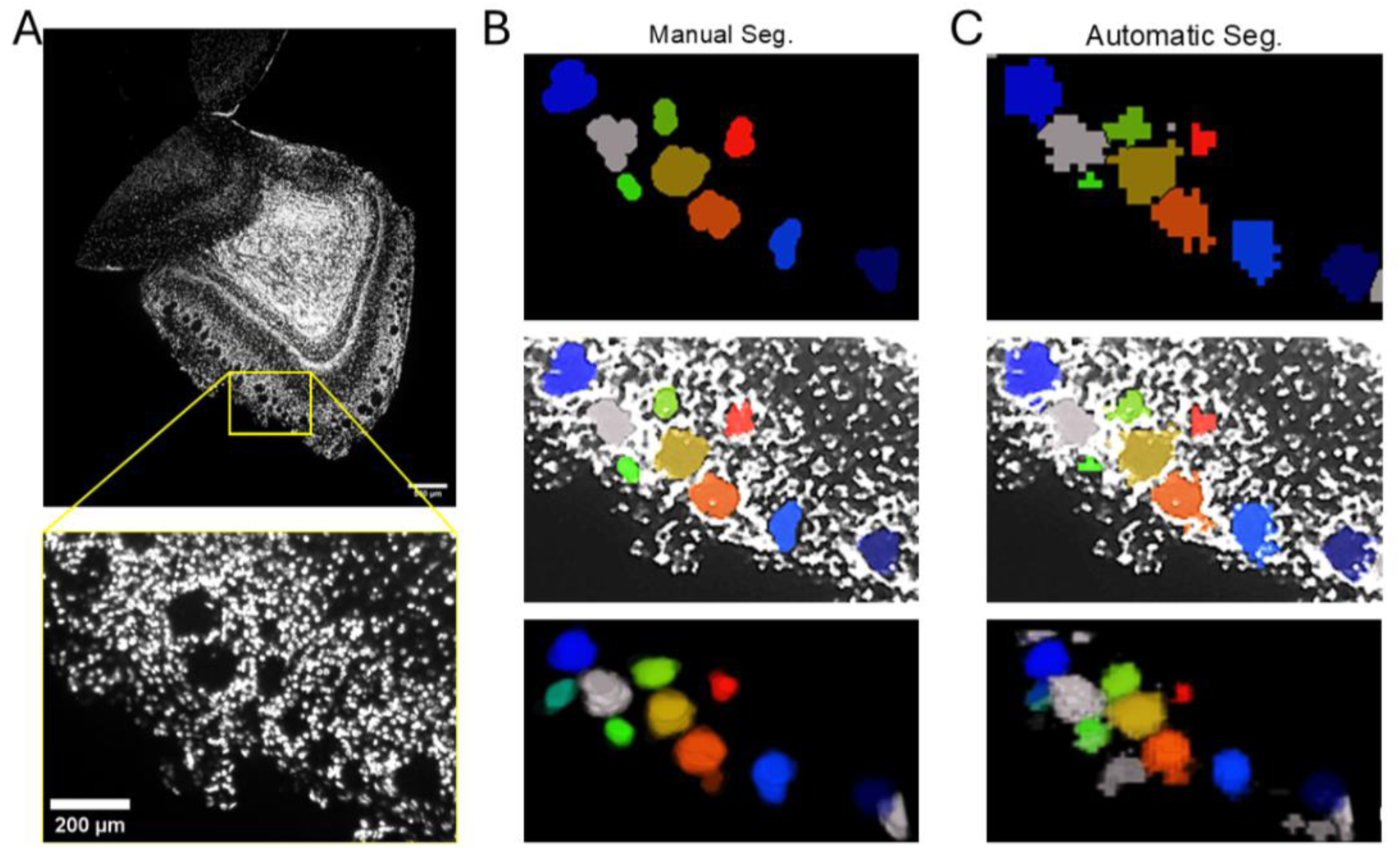
Manual three-dimensional segmentation validates automated reconstruction of OB glomeruli. (A) Representative optical section of an expanded mouse OB imaged by LSFEM using the Hoechst channel. The boxed region indicates the region of interest (ROI) selected for manual and automated three-dimensional analysis, shown at higher magnification on the right. (B) Manual segmentation of glomeruli within the selected ROI. Individual manually identified glomeruli are displayed using arbitrary colors (middle-up), overlaid onto the corresponding LSFEM image (middle-center), and reconstructed across consecutive optical sections to visualize individual objects in three dimensions (middle-bottom). (C) Corresponding automated segmentation of the same ROI using the SegFormer-based segmentation, morphological post-processing, and three-dimensional reconstruction workflow. Individual automatically reconstructed glomeruli are displayed using arbitrary colors (right-up), overlaid onto the corresponding LSFEM image (right-middle), and visualized as three-dimensional objects (right-bottom).

### Glomerular candidates were reconstructed as three-dimensional objects across the complete OB volume

After morphological separation, we assembled consecutive processed masks in their original z-order and subjected them to three-dimensional connected-component analysis. This step associated corresponding glomerular candidates across neighboring optical sections, allowing individual glomeruli to be represented as reconstructed three-dimensional objects throughout the complete OB volume (Figure 7). The resulting reconstruction reproduced the characteristic peripheral organization of the glomerular layer and enabled visualization of individual reconstructed objects independently (Figure 7B) or in their anatomical context within the original LSFEM volume (Figure 7C). Most notably, this transformation from slice-wise segmentation to a whole-volume object representation provided the basis for subsequent automated quantification of glomerular number across complete OB datasets.

**Figure 7.**
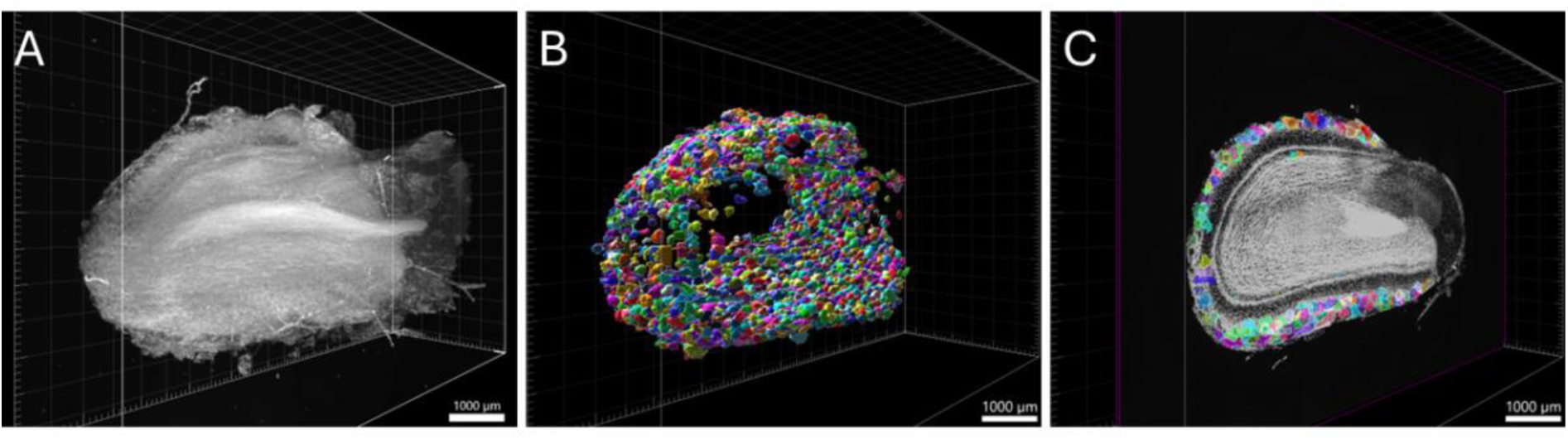
Three-dimensional reconstruction of segmented glomeruli across the OB. (A) Three-dimensional rendering of the complete expanded OB imaged by LSFEM using Hoechst nuclear staining. (B) Three-dimensional reconstruction of individual glomerular objects obtained following SegFormer-based segmentation, morphological post-processing, and connected-component analysis. Individual glomeruli are displayed using arbitrary colors for visualization. (C) Overlay of the reconstructed glomerular objects onto the corresponding LSFEM volume, illustrating their spatial distribution within the peripheral glomerular layer of the OB.

### Automated three-dimensional analysis enabled bulb-wide quantification of glomeruli

Having established the segmentation and three-dimensional reconstruction workflow, we next applied the complete analysis pipeline to 15 LSFEM OB datasets. Automated three-dimensional connected-component analysis identified individual reconstructed glomerular objects throughout each analyzed volume. Glomerular counts varied across datasets, with intact OB volumes generally yielding counts of approximately 2,000–3,000 glomeruli (Figure 8). In contrast, datasets in which the OB was physically damaged or incomplete showed substantially lower counts, emphasizing the importance of sample integrity for bulb-wide quantification. After excluding the three damaged datasets, the automated analysis yielded a median of 2,677 glomeruli per OB (Figure 8) (IQR, 2,154.5–2,899; range, 1,982–2,960) across the remaining 12 intact datasets. Variation in counts across datasets reflects differences in biological samples and imaging volumes, while consistent application of the same automated workflow enabled reproducible bulb-wide estimation of glomerular number across intact LSFEM datasets.

**Figure 8.**
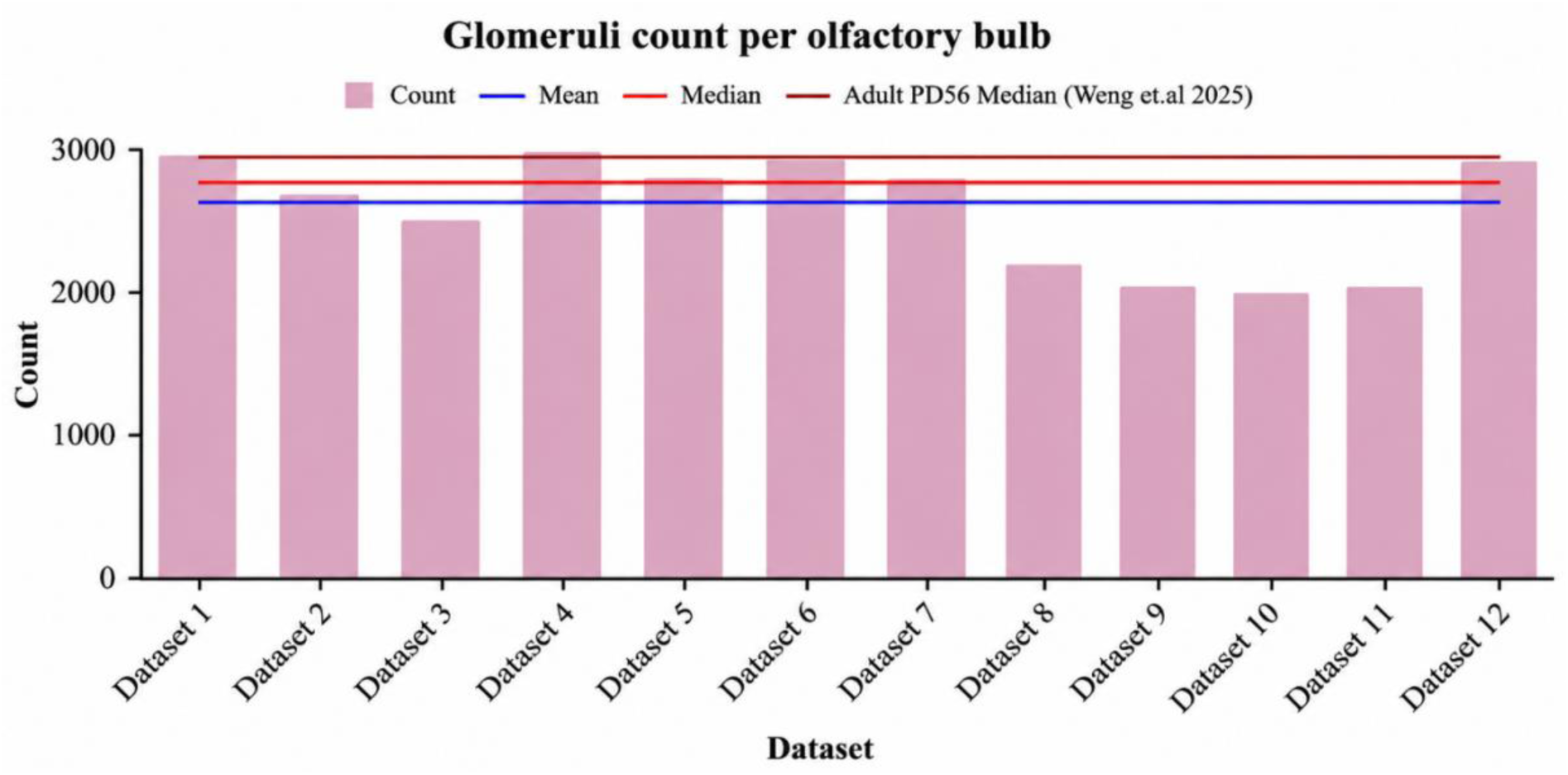
Quantitative summary. Automated Glomerular count across 12 OBs, with mean, median and PD56 median from Weng et al. (2025)

### Bulb-wide estimates approached the scale of an empirical anatomical evaluation

Finally, we placed the automated estimates in the context of existing anatomical measurements. The median count of 2,677 glomeruli was 6% below the adult PD56 median of 2,851 obtained by complete manual three-dimensional segmentation (Weng et al., 2025) but remained within the broad range reported across historical studies. Because the datasets differed in tissue source, labeling, sampling, and image processing, this comparison provides a scale reference rather than a direct accuracy test. The lower automated estimate is consistent with the cumulative effects of conservative foreground detection, reduced axial and lateral sampling, watershed partitioning, and connectivity-based merging. Importantly, the pipeline recovered the expected order of magnitude across all bulbs and converted large LSFEM volumes into reproducible, anatomically interpretable glomerulus segmentations.

### Glomeruli of different sizes are intermingled throughout the OB

Beyond determining total glomerular number, three-dimensional reconstruction allowed us to examine whether variation in glomerular morphology follows a spatial organization across the complete OB. For each reconstructed glomerulus, volume, sphericity, and three-dimensional centroid coordinates were extracted, providing a bulb-wide representation of individual morphological and spatial properties (Figure 9A, B). Glomeruli showed substantial variation in volume, yet mapping this variation back onto the reconstructed bulb revealed that small, intermediate, and large glomeruli were broadly intermingled throughout the glomerular layer, with no evident segregation into size-defined anatomical domains (Figure 9B). Classification based on volume and sphericity similarly showed that the resulting morphological groups were distributed across the OB rather than confined to discrete regions (Figure 9C, D). Thus, glomerular size heterogeneity emerges as a bulb-wide organizational feature. Despite pronounced differences between individual glomeruli, these differences do not translate into an obvious large-scale spatial partitioning of the glomerular population.

**Figure 9.**
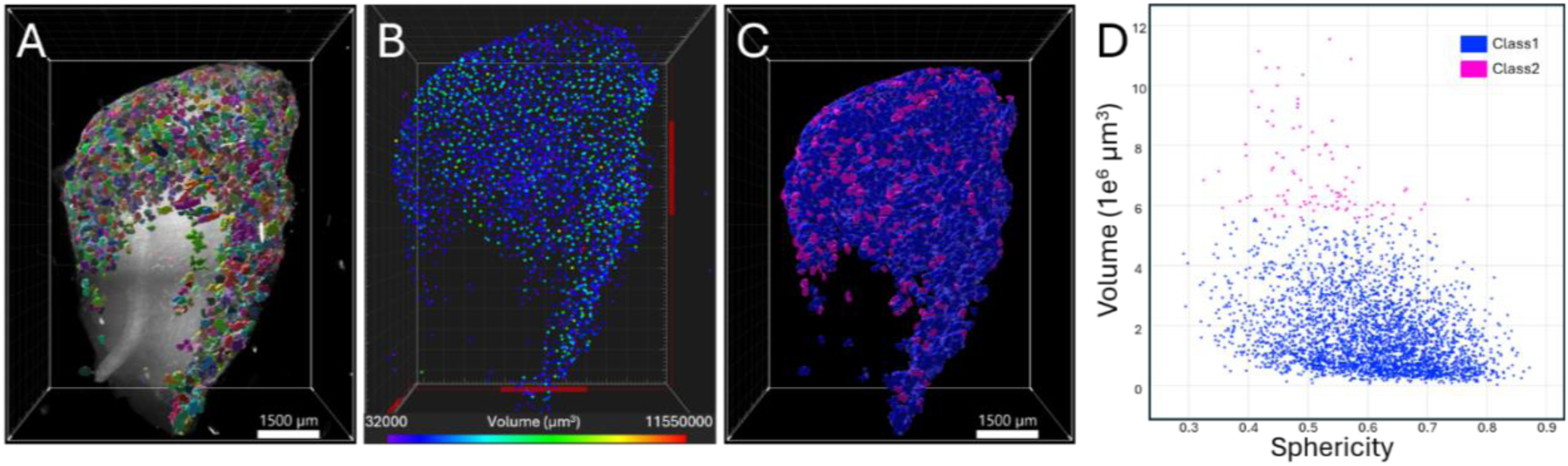
Three-dimensional reconstruction enables quantitative analysis of glomerular morphology across the OB. (A) Three-dimensional representation of a fully segmented OB, with individual reconstructed glomeruli displayed using arbitrary colors to illustrate their spatial distribution and individual object identities. (B) Centroid representation of the reconstructed glomeruli, in which each point corresponds to an individual glomerulus and is color-coded according to its reconstructed volume. (C) Three-dimensional visualization of glomeruli classified by morphological properties (volume and sphericity), illustrating the spatial distribution of glomeruli in the resulting morphological classes. (D) Relationship between reconstructed glomerular volume and sphericity for individual glomeruli, with colors corresponding to the classes displayed in (C). Class 1 glomeruli (blue) are characterized by lower volume and higher sphericity, whereas Class 2 glomeruli (magenta) show higher volume and lower sphericity

### Object-level reconstruction provides a relative spatial reference framework within the olfactory bulb

Because each reconstructed glomerulus retains a three-dimensional centroid coordinate, the object-level map can also be used to define relative spatial relationships within the intact OB. We first represented the position of all reconstructed glomeruli relative to the origin of the image volume (Figure 10A). Importantly, the reference position can be reassigned to any selected glomerulus or anatomically defined region within an individual bulb (Figure 10B), allowing the positions of all other reconstructed objects to be expressed relative to that location (Figure 10C). This does not require the selected glomerulus to represent the same receptor-defined glomerulus across different animals. Rather, it establishes a specimen-specific spatial framework in which additional anatomical signals, such as axonal projection or termination fields, can be related to the local organization of the glomerular layer. Such relative representations provide a basis for comparing where additional signals are positioned within the glomerular architecture of different OBs.

**Figure 10.**
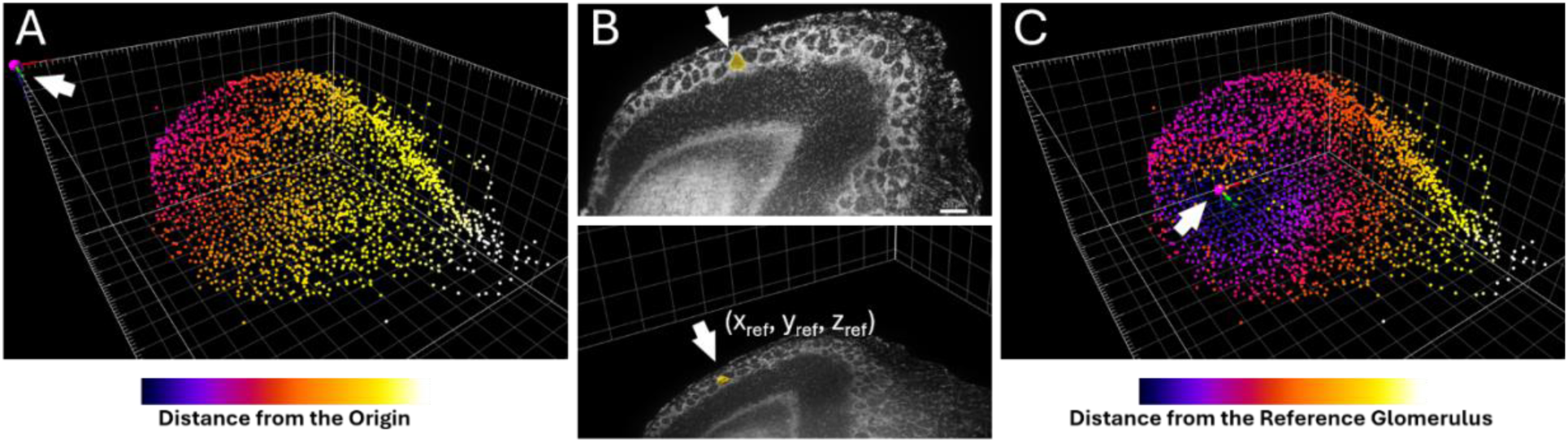
Reconstructed glomeruli provide a relative three-dimensional spatial reference framework within the OB. (A) Three-dimensional centroid representation of individually reconstructed glomeruli. Each point corresponds to the centroid of a segmented glomerular object and is color-coded according to its Euclidean distance from the origin of the image volume. The color scale represents increasing distance from this reference position. (B) Example illustrating how an individual reconstructed glomerulus can be selected as a local reference point (x_ref_, y_ref_, z_ref_) within a given OB. The selected glomerulus is highlighted in yellow and indicated by the white arrow in the corresponding LSFEM image and three-dimensional reconstruction. (C) The centroid of the selected glomerulus is assigned as the new reference coordinate, and the relative distance of every reconstructed glomerulus from this position is calculated. This analysis illustrates how object-level reconstruction can provide a specimen-specific spatial framework for relating additional anatomical signals to the local organization of the glomerular layer.

## Discussion

### Integrating whole-bulb LSFEM with automated reconstruction

This study establishes an end-to-end experimental and computational framework for reconstructing thousands of individual glomeruli across intact mouse OBs. A key advantage of our approach is the integration of whole-organ LSFEM with automated object-level analysis. Expansion and light-sheet fluorescence microscopy enable imaging of the entire OB from one hemisphere without physical sectioning, preserving its continuous three-dimensional architecture while retaining complementary cellular information from nuclear staining, genetically encoded fluorescence, and post-expansion immunolabeling. Each reconstructed glomerulus can therefore be retained within the anatomical context of the intact bulb rather than being analyzed as an isolated profile or within a restricted tissue region.

The computational workflow transforms this continuous volumetric information into a bulb-wide map of individual glomerular objects. SegFormer identifies the glomerular foreground from image context (Figures 3 and 4), morphology-based post-processing separates closely apposed glomeruli that remain connected in the semantic predictions (Figure 5), and three-dimensional connected-component analysis links the resulting candidates across consecutive optical sections. The methodological advance therefore lies not in a single segmentation step, but in the continuous sequence from intact whole-bulb imaging to object-level reconstruction and quantification. By combining these components, complex LSFEM volumes containing thousands of glomeruli are converted into quantitative three-dimensional maps in which individual objects retain their morphology and spatial position within the complete OB.

### From glomerular field segmentation to individual glomeruli

Our results illustrate why glomerular quantification cannot be reduced to a conventional foreground-segmentation problem. SegFormer reliably identified the glomerular layer across variations in OB geometry and signal distribution, yet neighboring glomeruli frequently remained connected within the predicted masks (Figure 4). This distinction between anatomical localization and object identity is also reflected in the segmentation metrics (Table 1). High specificity and MCC indicate robust background rejection and strong overall agreement with the ground-truth masks, whereas the more moderate precision, recall, and Dice values reflect remaining differences between predicted and ground-truth masks, particularly at glomerular boundaries and within densely packed regions. Thus, semantic segmentation provides a reliable representation of the glomerular field, but an additional level of analysis is required to transform this representation into individual anatomical objects.

Morphological postprocessing provides this transition from semantic to object-level information. Iterative H-maxima detection and watershed segmentation exploit internal mask geometry to divide continuous foreground regions into discrete glomerular candidates (Figure 5). This strategy is particularly suited to the densely packed glomerular layer because it does not require visible gaps between adjacent structures. At the same time, the resulting object definition necessarily depends on glomerular geometry. The tuberiform, elongated, and irregular morphologies described by Weng and colleagues (Weng et al., 2025) can complicate the separation of closely apposed structures, generating an inherent balance between over-separation and residual merging. The resulting count should therefore be interpreted as the output of an explicit geometric definition of object identity rather than as a direct readout of the semantic mask. This transition from semantic foreground to individual object identity is essential for whole-bulb reconstruction. By applying the same geometric criteria throughout each volume, thousands of glomerular profiles can be converted reproducibly into discrete anatomical objects and compared using a common analytical definition across complete bulbs and across animals.

### Three-dimensional reconstruction and interpretation of glomerular counts

The transition from 2D segmentation to 3D objects introduces a second geometric criterion. Using 18-connectivity, voxels connected through faces or edges are assigned to the same object across neighboring optical sections, while structures touching only at their corners remain separated. This provides a practical compromise between fragmenting continuous glomeruli and unintentionally merging neighboring structures. Consequently, 3D object identity reflects not only the biological continuity of a glomerulus, but also the axial sampling of the image volume and the connectivity criterion applied during reconstruction.

The three-dimensional reconstruction was supported by two complementary validation levels. A synthetic repeated-slice control confirmed the expected behavior of axial linking under idealized conditions (Supplementary Figure S2), whereas manual three-dimensional segmentation of a real LSFEM volume provided the more stringent validation of the complete reconstruction workflow under biologically realistic conditions (Figure 6). Within this volume, manual reconstruction identified 28 individual glomeruli and the automated workflow identified 27. This close correspondence is particularly relevant because the comparison incorporates the natural changes in glomerular shape, intensity, and appearance that occur across consecutive optical sections. The result therefore supports the workflow’s ability to transform serial glomerular profiles into individual three-dimensional objects while preserving their local spatial organization.

At the whole-bulb level, the automated reconstruction yielded glomerular numbers that closely approached the anatomical scale obtained by complete manual three-dimensional reconstruction. Across the 12 intact datasets, the median automated count was 2,677 glomeruli per bulb, only 6% below the median of 2,851 reported by Weng et al. (2025) (Figure 8). This numerical proximity is notable given that the two studies used different tissue preparations, labeling strategies, imaging approaches, sampling parameters, ages, and strains and therefore do not constitute a matched accuracy comparison. Several methodological factors could contribute to the lower automated estimate, including the 10 µm axial sampling interval, image resizing, omission of weak foreground during segmentation, residual merging following morphological separation, and persistence of narrow connections between neighboring structures during three-dimensional reconstruction. The comparison should therefore be interpreted as an anatomical benchmark rather than a direct validation against the Weng dataset (Weng et al., 2025). Nevertheless, recovering thousands of individual objects and a bulb-wide count within approximately 6% of a complete manual reconstruction supports the anatomical plausibility of the automated reconstruction and demonstrates that the workflow captures the expected scale of the glomerular population.

### Scalable whole-bulb reconstruction enables comparative anatomy

A major practical consequence of the end-to-end workflow is that whole-bulb reconstruction becomes scalable. Complete manual three-dimensional segmentation has been reported to require approximately 90 h per OB (Weng et al., 2025), whereas computational processing of a complete dataset with the present workflow required approximately 9 h with substantially reduced manual intervention. This represents an approximately ten-fold reduction in analysis time. Combined with rapid whole-bulb LSFEM acquisition, automated segmentation and reconstruction therefore shift glomerular mapping from a labor-intensive reconstruction of individual specimens toward an analysis that can be applied systematically across multiple complete bulbs.

Application of the workflow across the present collection of independently acquired LSFEM volumes demonstrates the feasibility of repeated whole-bulb analysis, while also identifying tissue integrity as an important prerequisite for quantitative comparison. This scalability changes the type of anatomical questions that can be addressed. Glomerular architecture can be treated as a quantitative phenotype and compared across animals, developmental stages, genetic backgrounds, experimental conditions, or disease models while maintaining the same analytical criteria for every specimen. Bulb-wide differences in glomerular number, morphology, spatial distribution, or local organization can therefore be addressed at the population level rather than being restricted to detailed reconstruction of individual specimens. The combination of rapid volumetric imaging and automated reconstruction thus provides a route from descriptive whole-organ anatomy toward comparative and statistically powered analysis of OB architecture.

### Object-level reconstruction reveals bulb-wide morphology and supports multichannel mapping

Reconstructing glomeruli as individual three-dimensional objects also extends the analysis beyond total number. Because every reconstructed glomerulus retains its identity, morphology, and spatial coordinates, properties such as volume and sphericity can be mapped directly back onto the intact OB. Moreover, centroid coordinates allow individual glomeruli and additional anatomical signals to be expressed relative to defined positions within each reconstructed bulb, providing a specimen-specific spatial reference framework (Figure 10). This analysis revealed substantial variation in glomerular volume, yet small, intermediate, and large glomeruli were broadly intermingled throughout the glomerular layer, without evident segregation into size - defined anatomical domains (Figure 9). Classification based on volume and sphericity produced a similar pattern, with morphological groups distributed across the reconstructed bulb rather than confined to discrete regions. Thus, pronounced morphological heterogeneity among individual glomeruli does not translate into an obvious large-scale partitioning of the OB according to glomerular size.

A particularly useful consequence of this object-based representation is that spatial relationships can be expressed relative to the glomerular architecture itself rather than solely in microscope coordinates. The selected reference position does not need to correspond to the same receptor-defined glomerulus across animals. Instead, each OB can be analyzed within its own reconstructed anatomical framework, and the location of additional signals can be described relative to local glomerular organization, broader anatomical regions, or normalized bulb coordinates. This is particularly relevant for projection anatomy, where axonal termination fields can be mapped onto the reconstructed glomerular layer, and their relative position, extent, and regional distribution can be compared across specimens.

Importantly, the absence of obvious bulb-wide segregation by size alone does not exclude regional specialization of defined glomerular subpopulations. Previous work in the rat OB identified glomerular classes with distinct serotonergic and cholinergic innervation patterns that were also associated with differences in size, morphology, and topographic position (Gómez et al., 2005). These findings underscore that morphological properties may acquire greater biological meaning when considered together with molecular identity, connectivity, or modulatory input rather than as isolated parameters. This biological observation illustrates the additional information gained when thousands of glomeruli are retained as individually identifiable objects within a common whole-bulb coordinate space.

A second important feature of the approach is that whole-bulb reconstruction is embedded within a multichannel imaging framework rather than being restricted to a single anatomical signal. In the present datasets, Hoechst nuclear staining was combined with preserved genetically encoded tdTomato fluorescence in mitral and tufted cells, which could additionally be amplified by post-expansion immunolabeling. Neurofilament-H immunostaining further demonstrated that conventional antibody labeling can be introduced into the intact expanded preparation and visualized across the three-dimensional tissue volume (Supplementary Figure 1). Thus, structural, genetically encoded, and immunolabeled signals can be acquired within the same intact sample and registered to the same anatomical volume in which individual glomeruli are reconstructed. This creates a whole-bulb reference frame in which glomerular architecture can be related directly to additional cellular and molecular features rather than analyzed in isolation.

The object-based three-dimensional reference frame substantially expands the types of relationships that can be examined using the approach. Depending on the biological question, additional fluorescence channels can identify defined neuronal populations, axonal projections, interneurons, glial cells, vascular structures, or other molecular features and relate their spatial organization directly to the reconstructed glomerular map. For axonal projections, this would allow termination fields to be assigned to defined regions of the glomerular map and compared across animals based on their relative position and spatial extent, without requiring identification of homologous individual glomeruli. Genetically labeled mitral and tufted cell populations, for example, provide a route for examining how glomerular input architecture relates to downstream projection neurons within the same intact bulb. The same anatomical framework could also be combined with prior *in vivo* physiological measurements (Le-Jahns et al., 2026), including virally expressed calcium indicators or recording-site and probe-tract reconstruction, to register functionally characterized signals onto the subsequently reconstructed whole-bulb anatomy. The significance of multichannel LSFEM is therefore not simply the addition of further labels, but the possibility of integrating glomerular, cellular, molecular, and ultimately functional information within a common three-dimensional representation of the intact OB.

Taken together, the central advance of this study is an end-to-end route from intact whole-bulb imaging to scalable three-dimensional reconstruction of thousands of individual glomeruli. By integrating LSFEM with automated semantic segmentation, instance separation, and three-dimensional object reconstruction, complex whole-organ image volumes are converted into quantitative anatomical maps while preserving the position and morphology of each reconstructed object. The workflow produced bulb-wide glomerular numbers approaching those obtained by complete manual reconstruction while reducing analysis time by approximately an order of magnitude. At the same time, object-level mapping revealed that glomerular size heterogeneity is distributed across the bulb without obvious large-scale size-defined domains. Because LSFEM retains genetically encoded fluorescence and supports post-expansion immunolabeling, these reconstructed glomerular maps can further serve as a common three-dimensional reference for additional cellular and molecular information. Together, these capabilities move whole-bulb glomerular mapping from labor-intensive reconstruction of individual specimens toward scalable, multimodal analysis of OB architecture.

## Resource availability

Trained model weights were saved locally. Repository links for raw data, manual masks, source code, trained weights, and analysis outputs are available upon request.

## Author contributions

| Author | Term | Definition |
|---|---|---|
| JER-G, MKS | Conceptualization | Ideas; formulation or evolution of overarching research goals and aims |
| JER-G, JIJ, SIG, AM, MKS | Methodology | Development or design of methodology; creation of models |
| JER-G, JIJ, SIG, AM | Formal analysis | Application of statistical, mathematical, computational, or other formal techniques to analyze or synthesize study data |
| JER-G, JIJ, SIG, DR | Investigation | Conducting a research and investigation process, specifically performing the experiments, or data/evidence collection |
| TA, MKS | Resources | Provision of study materials, reagents, materials, patients, laboratory samples, animals, instrumentation, computing resources, or other analysis tools |
| JER-G, MKS | Writing - Original Draft | Preparation, creation and/or presentation of the published work, specifically writing the initial draft (including substantive translation) |
| JER-G, JIJ, SIG, DR, AM, TA, MR, MKS | Writing - Review & Editing | Preparation, creation and/or presentation of the published work by those from the original research group, specifically critical review, commentary or revision – including pre-or postpublication stages |
| JER-G, JIJ, SIG | Visualization | Preparation, creation and/or presentation of the published work, specifically visualization/ data presentation |
| JER-G, TA, MR, MKS | Supervision | Oversight and leadership responsibility for the research activity planning and execution, including mentorship external to the core team |
| JER-G, MKS | Project administration | Management and coordination responsibility for the research activity planning and execution |
| TA, MKS | Funding acquisition | Acquisition of the financial support for the project leading to this publication |

## Acknowledgements

We thank Lydia Fischer for excellent technical support. M.K.S. was supported by German Research Foundation (SFB1089 P02, P03 and B06; SPP 2041 SCHW1578/2-1 and SPP 2411 SCHW2100/1-1) and the program Netzwerke 2021, an initiative of the Ministry of Culture and Science of the State of Northrhine Westphalia.

MR was supported by DFG 445965029/RO4046/5-1, 466488864/RO4046/6-1 466488864/RO4046/7-1, 520287065/RO4046/8-1 and Research Training Group 362321501/GRK2413.This work was further supported by the German Research Foundation (FOR5424 “Modolfor”, M.R. and T.A) and the European Union (ERC, “TempCOdE”, 101077017, T.A.).

This work was also supported by priority program SPP 2411, project number 520617944 (M.K.S. and M.R.)

Views and opinions expressed are, however, those of the author(s) only and do not necessarily reflect those of the European Union or the European Research Council. Neither the European Union nor the granting authority can be held responsible for them.

