## supplementary figures for "Whole-bulb LSFEM enables automated reconstruction and quantitative spatial mapping of thousands of olfactory glomeruli"

**Supplements_V16**

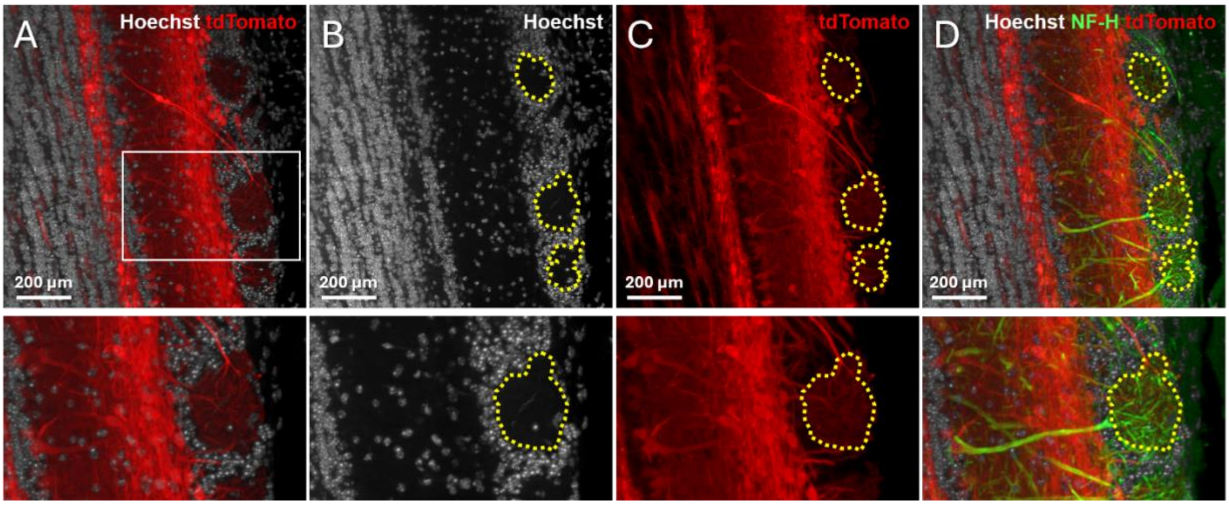


**Figure S1. Multichannel LSFEM visualization of glomerular architecture in the expanded mouse olfactory bulb.**
(A) Representative maximum-intensity projection (MIP; 20 optical sections) of an expanded mouse OB showing Hoechst 33342 nuclear staining (grey) and tdTomato-labelled mitral/tufted cells and their processes (red). The boxed region indicates the region of interest (ROI) shown at higher magnification below. (B) Hoechst channel of the same field. Putative glomeruli are visible as relatively cell-poor regions within the surrounding densely nucleated tissue and are indicated by yellow dotted outlines. The enlarged ROI is shown below. (C) Corresponding tdTomato channel showing labelled mitral/tufted cell processes extending into the Hoechst-defined cell-poor regions, providing complementary anatomical information supporting their identification as glomeruli. (D) Multichannel overlay combining Hoechst (grey), tdTomato (red), and neurofilament-H (NF-H; green) immunolabelling. NF-H staining illustrates how additional molecular and structural information can be incorporated into the same expanded preparation and spatially related to identified glomeruli. Yellow dotted outlines correspond to the same representative glomerular regions indicated in (B) and (C). Scale bars, 200 µm.

### Synthetic-stack control of three-dimensional object linking

To evaluate the behavior of the three-dimensional linking procedure independently of natural variation between consecutive optical sections, we generated a synthetic image stack with fully controlled axial continuity. A representative LSFEM section containing 197 manually counted glomerular profiles was replicated across 100 consecutive z-planes, such that the expected number of three-dimensional objects remained 197 despite the presence of identical profiles in every section (Supplementary Figure S2A,B). The synthetic volume was processed using the same segmentation, morphological post-processing, and connected-component analysis applied to the whole-bulb datasets. The pipeline reconstructed 202 three-dimensional objects, corresponding to an absolute difference of five objects (2.5%) from the expected count (Supplementary Figure S2C,D). Importantly, replication of the profiles across 100 optical planes did not result in proportional overcounting, demonstrating that spatially continuous detections were linked across the z-axis rather than treated as independent objects in individual sections. This synthetic control therefore confirms the expected behavior of the axial linking procedure under idealized conditions of spatial continuity.

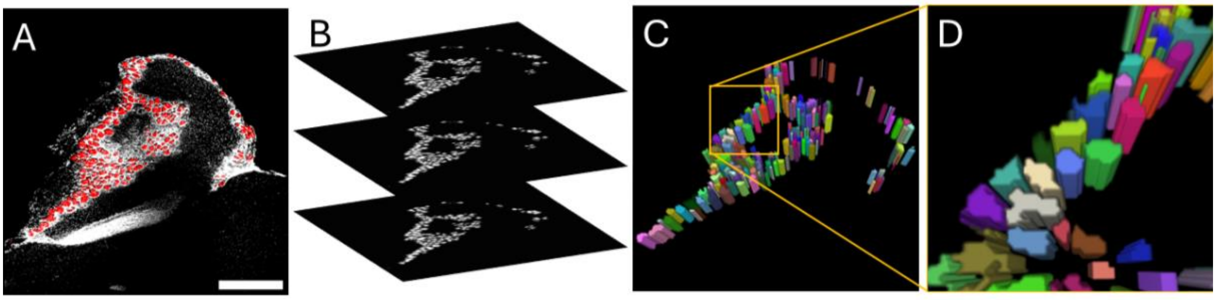


**Figure S2. Synthetic-stack control of three-dimensional glomerular linking.** (A) Representative 2D LSFEM image of the OB with the corresponding predicted glomerular mask overlaid in red. (B) Schematic representation of the synthetic volume generated by replicating the same 2D mask across 100 consecutive z-planes. (C) Three-dimensional reconstruction obtained from the synthetic stack following connected-component analysis, in which spatially connected regions across consecutive planes are linked to form individual 3D glomerular objects. Individual reconstructed objects are displayed using arbitrary colors for visualization. (D) Enlarged view of the boxed region in (C), illustrating the assignment of a distinct identity to each reconstructed 3D object.
